# Label-Free Mammalian Cell Tracking Enhanced by Precomputed Velocity Fields

**DOI:** 10.1101/2023.01.25.525598

**Authors:** Yue Han, Yang Lei, Viktor Shkolnikov, Daisy Xin, Steven Barcelo, Jan Allebach, Edward J. Delp

## Abstract

Label-free cell imaging, where the cell is not “labeled” or modified by fluorescent chemicals, is an important research area in the field of biology. It avoids altering the cell’s properties which typically happens in the process of chemical labeling. However, without the contrast enhancement from the label, the analysis of label-free imaging is more challenging than label-based imaging. In addition, it provides few human interpretable features, and thus needs machine learning approaches to help with the identification and tracking of specific cells. We are interested in label-free phase contrast imaging to track cells flowing in a cell sorting device where images are acquired at 500 frames/s. Existing Multiple Object Tracking (MOT) methods face four major challenges when used for tracking cells in a microfluidic sorting device: (i) most of the cells have large displacements between frames without any overlap; (ii) it is difficult to distinguish between cells as they are visually similar to each other; (iii) the velocities of cells vary with the location in the device; (iv) the appearance of cells may change as they move in and out of the focal plane of the imaging sensor that observes the isolation process. In this paper, we introduce a method for tracking cells in a predefined flow in the sorting device via phase contrast microscopy. Our proposed method is based on DeepSORT and YOLOv4 and exploits prior knowledge of a cell’s velocity to assist tracking. We modify the Kalman filter in DeepSORT to accommodate a non-constant velocity motion model and integrate a representative velocity field obtained from fluid dynamics into the Kalman filter. The experimental results show that our proposed method outperforms several MOT methods for tracking cells in the sorting device.

## I. Introduction

Label-free cell imaging is increasingly gaining interest in biomedical research, as chemical labeling processes risk altering the cells’ properties and should be avoided, especially if the cells are to be used further for clinical applications [1]. ^1^ However, label-free imaging is more challenging to be analyzed than traditional label-based imaging, such as fluorescent imaging, since the contrast between the cell and the image background is typically low [2], the signal-to-noise ratio is often low as well [3], and specific molecular markers for tracking are generally unavailable. Furthermore, label-free imaging provides few human-interpretable image features (HIFs) and thus can benefit from machine learning approaches to help with identification and tracking of specific cell types. HIFs include cell sizes and shapes, nucleus sizes and shapes, textures, and morphologies [4].

In this paper, we focus on phase contrast imaging of mammalian cells, which is a common type of label-free imaging technique [5]. We use phase contrast microscopy to track cells flowing in a microfluidic sorting device. (referred to as the “chip” in this paper). Microfluidics is increasingly used for high throughput cell manipulations, including for clinical applications [6], and thus label-free tracking of cells in a flow is rapidly gaining importance. Figure 1 shows a frame taken of the microfluidic cell sorting device or the “chip”. The chip operates by a hydrodynamically focusing stream of cells that enters the chip hardware from the bottom inlet. Dielectrophoretic force guides the cells to two different outlets based on the polarization of the cells in an alternating current electric field. Since the cells are in a suspension state (i.e., suspended in a fluid as opposed to attached to a surface), they exhibit even fewer HIFs relative to their adhered state (they are adhered to a surface) and are thus difficult to track.

**Fig. 1:**
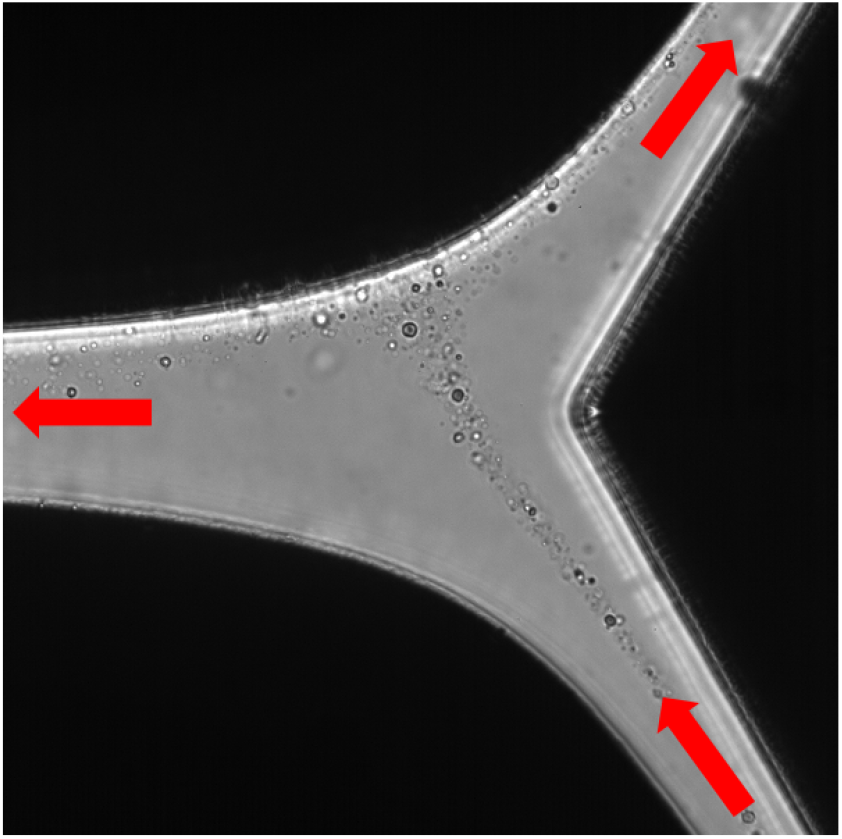
A phase contrast microscopy image of the cell sorting device or chip. The flow direction is shown in red arrows. A stream of cells enters the chip from the bottom inlet, the dielectrophoretic force guides the cells to two different outlets.

We introduce a method for tracking cells in a known flow field in the sorting device via phase contrast microscopy. We apply this method to track cells in our chip for the isolation of rare cells based on their electric properties. One major application of this system is the isolation of circulating tumor cells for therapy evaluation, where isolated cells are subject to therapy candidates and the cells’ response is tracked. We desire to use label-free sorting and cell tracking of these cells to avoid the risk of the chemical labeling or staining processes altering the cells’ response to therapy.

Furthermore, we require the sorting to be high throughput, since clinically relevant concentrations of circulating tumor cells can be as low as in the order of 10 cells in 10^10^ total blood cells. Besides the challenges of tracking the cells in label-free imaging, in our high speed imaging of the chip, we chose a frame rate based on a tradeoff between the need to track the cells for a longer time and camera data storage limitations. This results in a relatively large cell displacement between each frame (up to 6 to 10 cell diameters). This is a widely encountered tradeoff when imaging high throughput microfluidic systems with high speed cameras, which are often available to many microfluidic and biomedical researchers. Due to these challenges and constraints, existing multiple object tracking techniques do not perform well for tracking cells in the chip.

To address these challenges, we describe a new multiple object tracking (MOT) technique for label-free imaging assisted by a representative velocity field precomputed from the flow field in the chip. We present a simple and effective modification to the Kalman filter [7] to make full use of the representative velocity field. We first adjust the Kalman filter state matrix used in DeepSORT [8] to address the non-constant velocity motion of the cells in the chip. Then we modify the “measurement” part of the Kalman filter used in DeepSORT to integrate the representative velocity field into the motion estimation module.

These two modifications can be used with any MOT task when objects’ velocities are accessible before or during the tracking. Benefiting from these two modifications to the baseline DeepSORT method, improvements are achieved for both of the two commonly used metrics for MOT tasks, MOTA (Multiple Object Tracking Accuracy) [9] and IDF1 (Identification F1 Score) [10]. We tested our proposed method on our challenging cell tracking dataset and the evaluation results show our method outperforms several existing MOT methods and achieved 26.4 in MOTA and 34.7 in IDF1. Our major contributions are summarized as follows:

- We describe a cell tracking technique for tracking multiple cells in the chip.
- We propose modifications to the Kalman filter in DeepSORT to integrate velocity information of the tracked objects, which improves the tracking of multiple cells.

## II. Related Work

In this section, we describe the existing methods in general multiple object tracking, then we introduce some previous work that focuses on cell tracking.

### A. General Multiple Object Tracking

The goal of Multiple Object Tracking (MOT) is to analyze a video in order to identify and track objects that belong to one or more categories, such as pedestrians, cars, and animals without any prior knowledge about the appearance and the number of targets. The standard approach employed in MOT is tracking-by-detection [11], where a set of detections are extracted from the video frames and an association or correspondence method that assigns the same track identity number to the bounding boxes that contain the same object. The association methods rely on motion information or object appearance features (e.g., shape or edges), or both.

Motion estimation methods model how the object moves and predicts the position of the object in future frames. The MOT methods in [12]–[15] only used motion information to associate a track with a set of objects. Kalman filter [7] is commonly used for motion estimation in MOT [8], [15]–[17]. With the recent development of neural networks, the recurrent neural network (RNN) or long short-term memory (LSTM) network has been used for non-linear motion estimation in several MOT approaches [12], [18], [19]. However, these methods usually require a large amount of ground truth data (more than 5000 tracks) to train the neural network.

The object appearance features are unique visual representations of an object to distinguish it from other objects. Examples of appearance features include shapes, sizes, colors, and edges. Convolutional Neural Network (CNN) is commonly used to extract the visual features. MOT methods described in [8], [20], [21] utilized both motion information and appearance features for associating tracks with particular objects. The addition of the appearance features assists the motion estimation module to deal with the occlusion of objects and to alleviate the problem of the objects changing their identities, therefore achieving better tracking results than methods solely using the motion information.

With the latest progress in the object re-identification, several methods [22]–[25] only used the object appearance features for track association. However, these methods require the appearance features of an object to be similar across the frames, and have enough differences from other objects.

### B. Cell Tracking

Although deep learning methods have dominated the analysis of the MOT problem for natural images, there are only a few deep learning approaches for cell tracking [26], [27]. Unlike the general MOT, a very limited number of publicly available ground truth datasets focus on cell tracking. The ISBI Cell Tracking Challenge [28] had been the only standardized benchmark in this area. However, the evaluation methodology of this challenge is different from the common metrics generally used in MOT. CTMC Cell Tracking Challenge [29] is the first published cell tracking dataset with an online evaluation server using both evaluation metrics from mainstream MOT and cell tracking communities. The difference within cell tracking tasks, such as having distinct motion patterns and whether containing the divisions of parent cells, are significant. Unlike the methods designed for the challenges mentioned above that focus on tracking live-cells’ interaction with their surrounding environment, tracking cells in the chip involves hydrodynamic drag force which accelerates cell movement. This major difference brings unique challenges which cause other cell tracking methods not to work well on tracking cells in the chip.

## III. Proposed Method

Tracking cells in the chip is challenging because the cells have non-constant velocities and appear similar. Since the chip utilizes a uniform dielectrophoretic force field to separate the cells, we are able to estimate a velocity field in the chip. We call this the representative velocity field and it represents the estimated velocities of the cells in the chip. We describe how we get this representative velocity field in detail in Section IV-B. The ideal cell tracking method should utilize both cell motion and cell appearance features for better track association. It also needs to take advantage of the representative velocity field to deal with the unique motion pattern in the chip. Computational efficiency is another important factor when analyzing long image sequences in real applications. Therefore, we choose DeepSORT [8] with YOLOv4 [30] detector as our baseline method and incorporate prior knowledge of the cell’s velocity into the motion estimation module.

### A. Baseline Cell Tracking

Our baseline cell tracking method uses a tracking-by-detection approach. First, we detect all the cells in each frame and construct a bounding box around each cell. Then a cell tracker is used to associate tracks with particular cells (track association). YOLOv4 [30] is used as the cell detector in our baseline method. It is an efficient and powerful one-stage object detection and classification system which can produce detection results in real time. DeepSORT [8] is used for tracking the cells, which utilizes both bounding box parameters and appearance features of the detected cells to associate the cells to existing tracks. The block diagram of DeepSORT is shown in Figure 2.

**Fig. 2:**
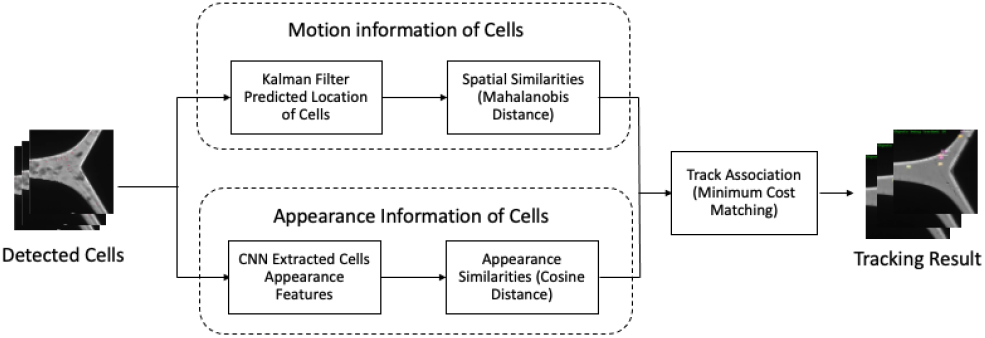
The block diagram of DeepSORT.

The motion information is estimated by using Kalman filter [7] that predicts the bounding box parameters of a cell in the next frame. The appearance features are determined using a pre-trained CNN through examining the pixels in the detected bounding box. The appearance features and the estimated motion are then used for matching a detected cell to an existing track. The Hungarian minimum cost method [31] is used for matching newly detected cells to previously tracked cells with similar motion and cell appearance.

DeepSORT [8] utilizes Kalman filter for tracking the bounding boxes of the cells. It provides a prediction of the future system state, based on the past estimation of the motion and the measurement of the newly detected cells’ motion. The system state vector x of the Kalman filter is defined as an 8-dimensional vector (*x*, *y*, *a*, *h*, *ẋ*, *ẏ*, *ȧ*, *ḣ*). Where (*x*,*y*) marks the center of the cell bounding box, *h* is the height of the bounding box, *a* is the aspect ratio of the bounding box (height/width), and (*ẋ*, *ẏ*, *ȧ*, *ḣ*) are their corresponding velocities. Since the cell detector does not provide velocity information, the velocities in the initial system state vector are manually chosen. The measurement vector **z** in Kalman filter represents a true system state **y** with random measurement noise. In DeepSORT, measurement vector **z** only contains cell’s bounding box information with velocities assumed 0. By incorporating the system state estimation **x_n_** calculated from the previous frame and the current measurement vector **z_n_**, the Kalman filter obtains refined current system state estimation 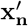, and uses state transition matrix F to predict the next system state x_n+1_ from 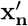. The state transition matrix **F** represents the system’s dynamic model, and DeepSORT uses a constant velocity motion model. The new prediction of the next system state x_n+1_ will be used as the system state estimation for further iterations.

The cell appearance feature in DeepSORT [8] is extracted by a CNN trained on a large-scale re-identification dataset. The appearance features of a detected cell are represented by a 1×128 vector. The distance between appearance vectors from the same cell across the frames should be small and the distance between appearance vectors from different cells should be large.

In DeepSORT, the cost *C* for associating two detected cells from two frames is defined in Equation 1

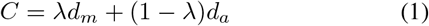

where *d_m_* is the Mahalanobis Distance [32] between the predicted system state and the measurement motion vector of the detected cell and *d_a_* is the Cosine Distance [33] between the appearance features of two cells. The weight λ is used to control how much the motion features or appearance features effect *C*. Pre-defined thresholds for the Mahalanobis Distance and the Cosine Distance are used to exclude outlier cases where two objects have significant differences in motion or appearance.

When associating detected cells with tracks, *C* is computed for each pair of an existing track and a detected cell in the new frame (removing outliers by thresholding). If *C* is the smallest between a detected cell and a confirmed track, then this cell is matched with the track and the unassociated age of this track is set to zero. The unassociated age of a track is the number of frames where a confirmed track fails to be associated with a detected cell. If a detected cell fails to associate with any of the confirmed tracks, then this cell is initialized as a new tentative track. For the detected cells in the following frame, we try to associate them with the new tentative tracks. If a new tentative track is successfully associated with a cell, it will be updated to a confirmed track. Otherwise, it will be removed immediately. For the confirmed tracks which fail to be associated with any newly detected cell in the frame, the unassociated age of this track will increase by 1. If the unassociated age exceeds the pre-defined maximum unassociated age, this track will be determined as ended.

### B. Integrating Velocity Prior Knowledge

Although DeepSORT has been widely used in many different tracking applications, it did not perform well in our dataset due to inaccurate motion estimation. Tracking cells in the chip requires more accurate motion estimation. The motion estimation method used in DeepSORT is a basic Kalman filter that assumes constant velocity which fails with non-constant velocity. Another issue with DeepSORT is that the velocities of newly detected cells are initialized to 0 because the detector cannot measure the velocity. The fact that cells move with different velocities and accelerations in the chip causes inaccurate motion estimation. In our application, we can get prior knowledge of cells’ velocities at different locations from the microfluidic chip design, which we call the representative velocity field. By incorporating these velocities into Kalman filter, we are likely to get better motion estimation.

#### 1) Modified Kalman Filter

To work with the non-constant velocity in the chip, we modify the Kalman filter state vector x to a 12-dimensional vector (*x*, *y*, *a*, *h*, *ẋ*, *ẏ*, *ȧ*, *ḣ*, *ẍ*, *ӱ*, *ä*, *ḧ*). Where (*x*, *y*) marks the center of the cell bounding box, h is the height of the bounding box, *a* is the aspect ratio of the bounding box (height/width), (*ẋ*, *ẏ*, *ȧ*, *ḣ*) are their corresponding velocities, and (*ẍ*, *ӱ*, *ä*, *ḧ*) are their corresponding accelerations.

With changes in the Kalman filter state vector, the state transition matrix **F** in the state extrapolation equation needs to be adjusted to a linear constant acceleration model. Equation 2 shows the extrapolation equation and Equation 3 shows matrix multiplication results.

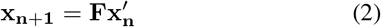

where *n* is the frame number, 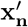 is the 12-dimensional refined system state estimation vector at frame n, x_n+1_ is the estimated next system state,and **F** is the modified transition matrix.

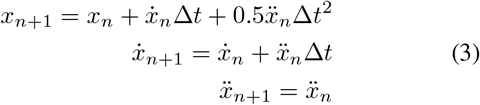

where *x* is the cell bounding box center position x-coordinate, *ẋ* and *ẍ* is its corresponding velocity and acceleration. Δ*t* is the time between two frames *n* and *n* + 1. The rest of the parameters in the system state vector are updated in the same way as *x*.

#### 2) Velocity Field Integration

To integrate the velocity field into the motion estimation module, we modify the “measurement” part of the Kalman filter. The measurement part takes the detected cell motion information and uses them to refine the Kalman filter estimated state vector. The measurement vector z used in the original Kalman filter is a 4-dimensional (*x, y, a, h*) vector. Where (*x, y*) are the cell bounding box coordinates, *h* is the height of the bounding box, and *a* is the aspect ratio of the bounding box (height/width). This 4-dimensional vector does not record any velocity information. We modify measurement vector z to a 8-dimensional (*x*, *y*, *a*, *h*, *ẋ*, *ẏ*, *ȧ*, *ḣ*) vector, where (*ẋ*, *ẏ*, *ȧ*, *ḣ*) are the corresponding velocities of the parameters in the original 4-dimensional measurement vector. In the measurement step, the (*x, y*) are obtained from the detected cell bounding box, we use the (*x, y*) location to find the corresponding velocities in the representative velocity field and use them for (*ẋ*, *ẏ*). The *ȧ* and *ḣ* are the rate of change of the aspect ratio and the height of the bounding box, we initialize them as 0 since the aspect ratio and the height of the cell do not change much.

With the new system state and measurement vectors, we adjust the observation matrix **H** to an 8×12 matrix with an 8×8 identity matrix concatenated with an 8×4 all zero matrix. The observation matrix **H** is used to identify which states are measured. For example, we are not able to measure the acceleration of the cell. This adjustment ensures the velocities of cells are passed into the measurement vector. Equation 4 shows our modification in detail.

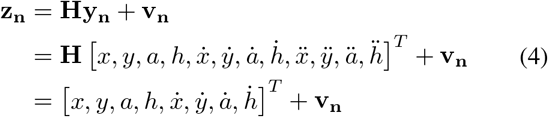

where **z_n_** is the measurement vector at frame n, **y_n_** is the newly measured true system state vector at frame n, **v_n_** is a Gaussian random noise vector, *x, y, a, h* are obtained from the cells’ bounding box, *ẋ* and *ẏ* are selected based on the location of the bounding box and the representative velocity field. The rest of the parameters are set to 0.

These modifications allow the Kalman filter motion estimation module in DeepSORT to take advantage of the velocities from the representative field and therefore model cells’ nonconstant velocities more accurately. The overall block diagram of our proposed cell tracking system is shown in Figure 4.

**Fig. 3:**
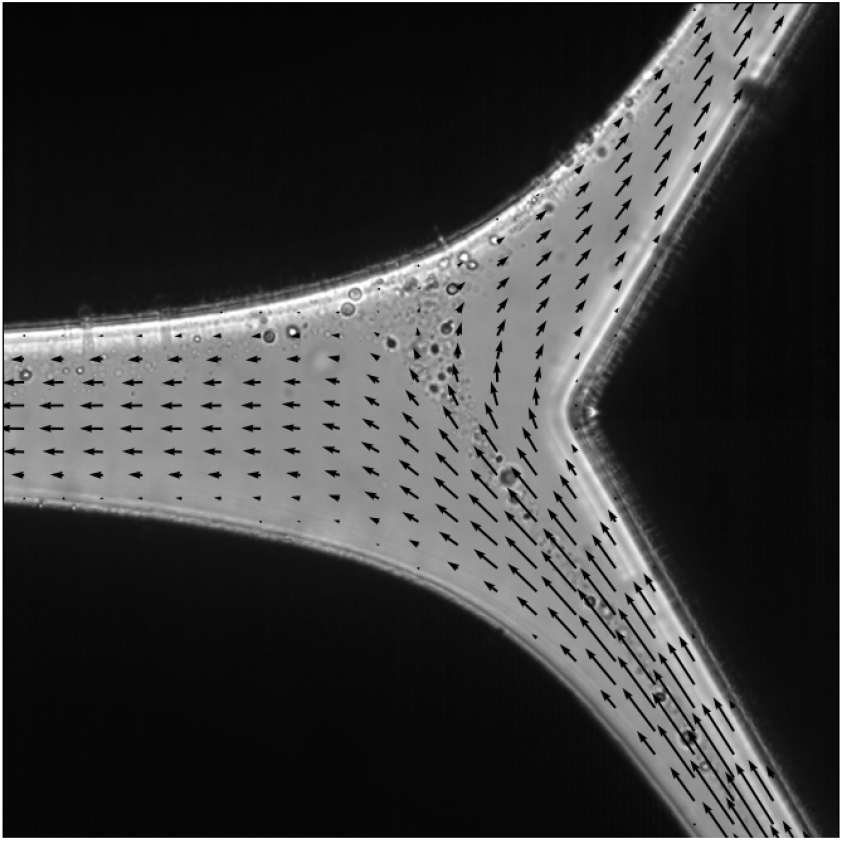
The representative velocity field in the chip. The velocity field in the chip is visualized as a collection of vectors with the magnitude of velocity and direction for that point.

**Fig. 4:**
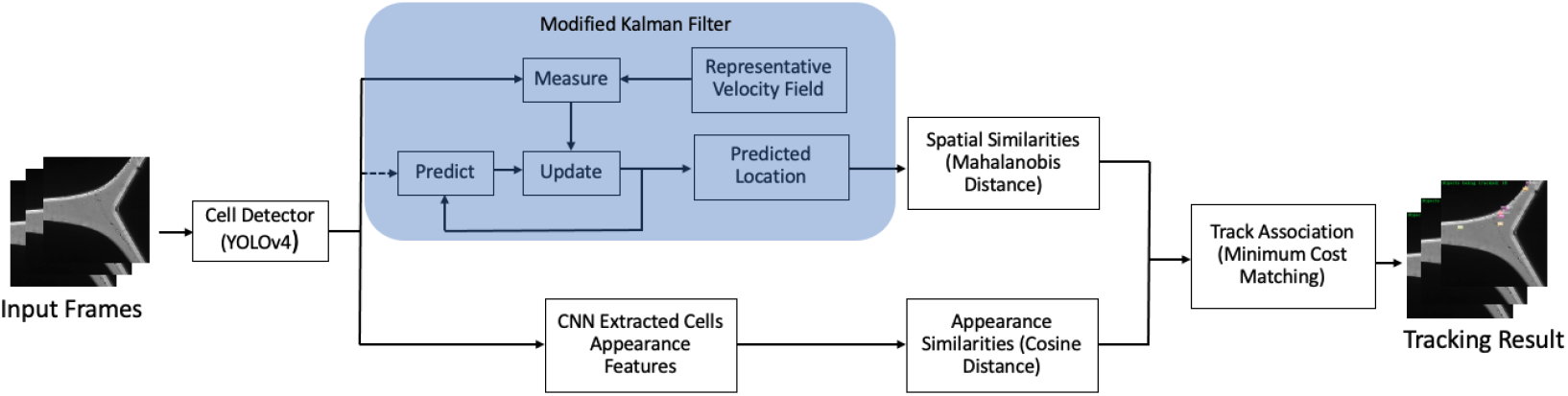
The block diagram of the proposed cell tracking system. (The dashed line process only happens once in initialization.)

## IV. Tracking Cells in the Chip

In this section, we introduce the type of data we acquired from our rare cell isolation chip. We present our cell tracking dataset and describe the challenges which make this cell tracking problem unique compared to general MOT problems and other cell tracking problems.

### A. Challenges

The video frames of the chip are phase contrast microscopy images captured by a high-speed camera attached to the microscope. The chip is designed to separate and isolate mixed cells, the high flow rate in this process causes large displacement of cells between the captured video frames. When cells move to the center of the chip, the dielectrophoretic force field slows them down for separation, then the cells accelerate when exiting the chip. Unlike most of the MOT tasks, large displacement of cells means there is no overlap of a cell’s bounding box between two frames. This unique motion pattern of the cells in the chip causes motion estimation methods to fail when trying to predict the location of the cells accurately.

The use of an object’s appearance features can help overcome the challenge of tracking objects with complicated motion patterns. It is also used to handle the occlusions in MOT. However, in our cell tracking problem, the difference in appearance between each cell is difficult to distinguish. The same individual cell may appear differently with a change of focus or location in the chip. Figure 5 shows the appearance of three different cells in 5 frames. This challenge demonstrates that appearance features alone are not enough for our cell tracking problem.

**Fig. 5:**
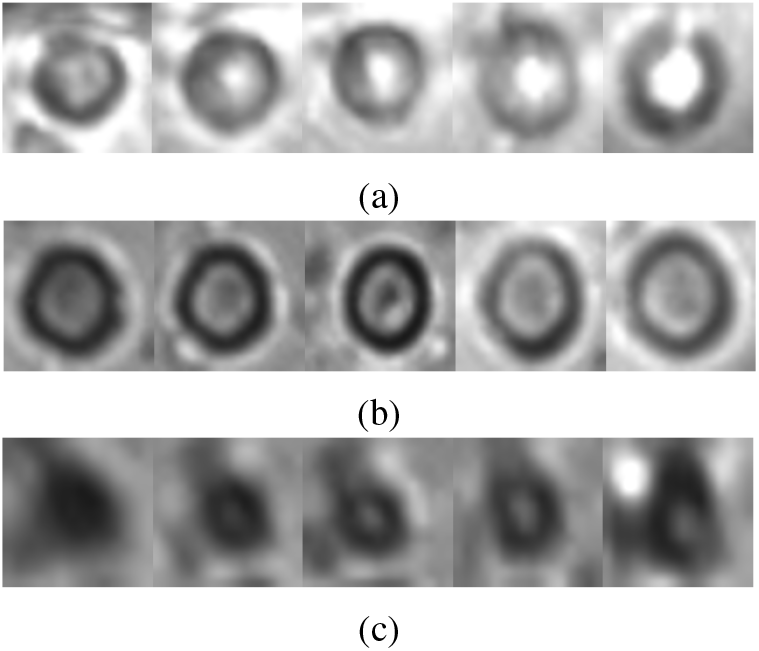
Each row is the appearance of a single cell in 5 different frames, (a)(b) are melanoma cancer cells, (c) is a white blood cell (WBC).

In summary, tracking cells in our chip consists of two main challenges: (i) the unique motion pattern of the objects with large displacements between frames, and (ii) the appearance of a cell is not distinct, and sometimes not consistent. These challenges exclude many well-known MOT methods for this problem.

### B. Representative Velocity Field

A cell’s velocity in the chip is determined by the superposition of the hydrodynamic drag force from the local flow velocity and a dielectrophoretic body force. In a large portion of the flow field, the contribution of the hydrodynamic drag force dominates. We thus used the flow velocity as the characteristic velocity field as shown Figure 3. We calculated this field by numerically solving the Navier-Stokes equations [34] using COMSOL [35]. We simplified the Navier-Stokes equations with a low Reynolds number assumption. For this solution, we prescribed the flow rate at the inlet and assumed equal pressures at the two outlets. We are able to estimate a cell’s velocity in the chip with this field information, and we shall denote this field as the “representative velocity field” of the chip.

The representative velocity field records horizontal and vertical velocities (*u_rx_*, *v_ry_*) at each location (*x, y*) in the chip. We mapped the velocity field to the image space by bilinear interpolation [36] to make sure every pixel in a frame has a representative velocity associated with it. The interpolation was done separately on each velocity component.

Although the representative velocity field can represent the general motion pattern of the cells in the chip, the cells’ velocities in each experiment may be different from the velocities obtained from the representative velocity field due to the different types of cells, the different flow rates used, the interactions between cells, and interaction with the chip boundaries. However, the Kalman filter can effectively produce motion estimation to predict the locations of the cells based on inaccurate and uncertain measurements. Thus, we use this representative velocity field as the prior information of a cell’s velocity to assist the tracking system to improve the cell tracking performance.

## V. Experimental Results

We conducted experiments using our own annotated/ground truth cell tracking dataset and compared our proposed method to the baseline DeepSORT and several other MOT methods. The tracking performance is evaluated with two commonly used metrics in MOT, which are Multi-Object Tracking Accuracy (MOTA) [9] and Identification F1 Score (IDF1) [10]. The score of MOTA ranges between negative values and 100 and IDF1 ranges between 0 and 100, both with better performance indicated by values lying closer to 100.

Our experiments were implemented on a 10-core Intel i9-9900X CPU@3.50GHz with an Nvidia TITAN RTX GPU. We selected the weight which controls the influence of motion or the appearance metric on the combined association cost λ = 0.6. YOLOv4 was trained for 500 epochs. The confidence threshold of the detector was set to 0.7 which means only the detected cells with a confidence score over 0.7 will be tracked.

### A. Dataset

A stream of melanoma cancer cells and white blood cells (WBC) mixture was injected into the input channel of the chip. The cell sorting process is captured by a high speed camera and phase contrast microscopy. The entire process contains more than 40,000 images captured at 500 frames per second. We randomly sampled 50 images to train the cell detector and excluded them from the dataset. We randomly selected the starting frame and annotated three 50-frame sequences. One of three 50-frame sequences is used for training the deep learning MOT methods for comparison. The other two are used for the evaluation of cell tracking methods. None of the three image sequences have any temporal overlap with each other. We are only interested in tracking in-focus cells, thus only the in-focus cells were annotated.

### B. Discussion

We compare our proposed method with the baseline DeepSORT [8] and two other popular MOT methods, FairMOT [25] and CenterTrack [37]. FairMOT [25] is a method which only uses object appearance features for detection and reidentification to get the tracking result. CenterTrack [37] localizes tracking objects and predicts their displacement with the previous frame to build the tracks. All these three methods showed top performance in the MOT challenge [38] for tracking pedestrians. We evaluate the comparison methods using the same dataset as used for our proposed method. The comparison results are shown in Table I. Due to limited amount of annotated training data and the challenges mentioned in section IV-A, the general MOT methods in our cell tracking task do not achieve decent results as they did in MOT challenges [38]. Our proposed method achieves higher MOTA and IDF1 results than all the comparison methods

**TABLE I:**
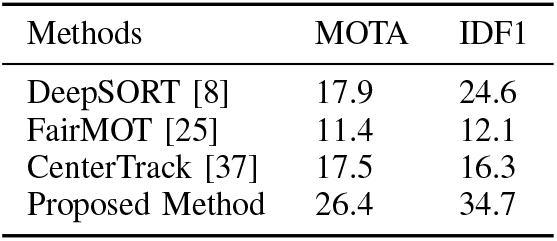
The cell tracking results of different methods

Figure 6 demonstrates our proposed method improves the cell tracking results in the chip by comparing it with the baseline method. By making use of the representative velocity field, our tracking method can successfully associate the tracks to the cells with high velocities. As shown in the third row in Figure 6, our method better capture cell 87 and cell 102 at the input area of the chip while these two cells did not show up in the baseline method results. Cell 7 from the third row of Figure 6 is a cell approaching the separation area, which means its motion changes from accelerating to slowing down. This cell is picked up for tracking sooner with our proposed method than the baseline method, which shows our method adapts better to the unique motion pattern in the chip. The rest of the results look similar, which means the deviation from real velocities in the representative field is well rectified by the modified KF and does not worsen the result.

**Fig. 6:**
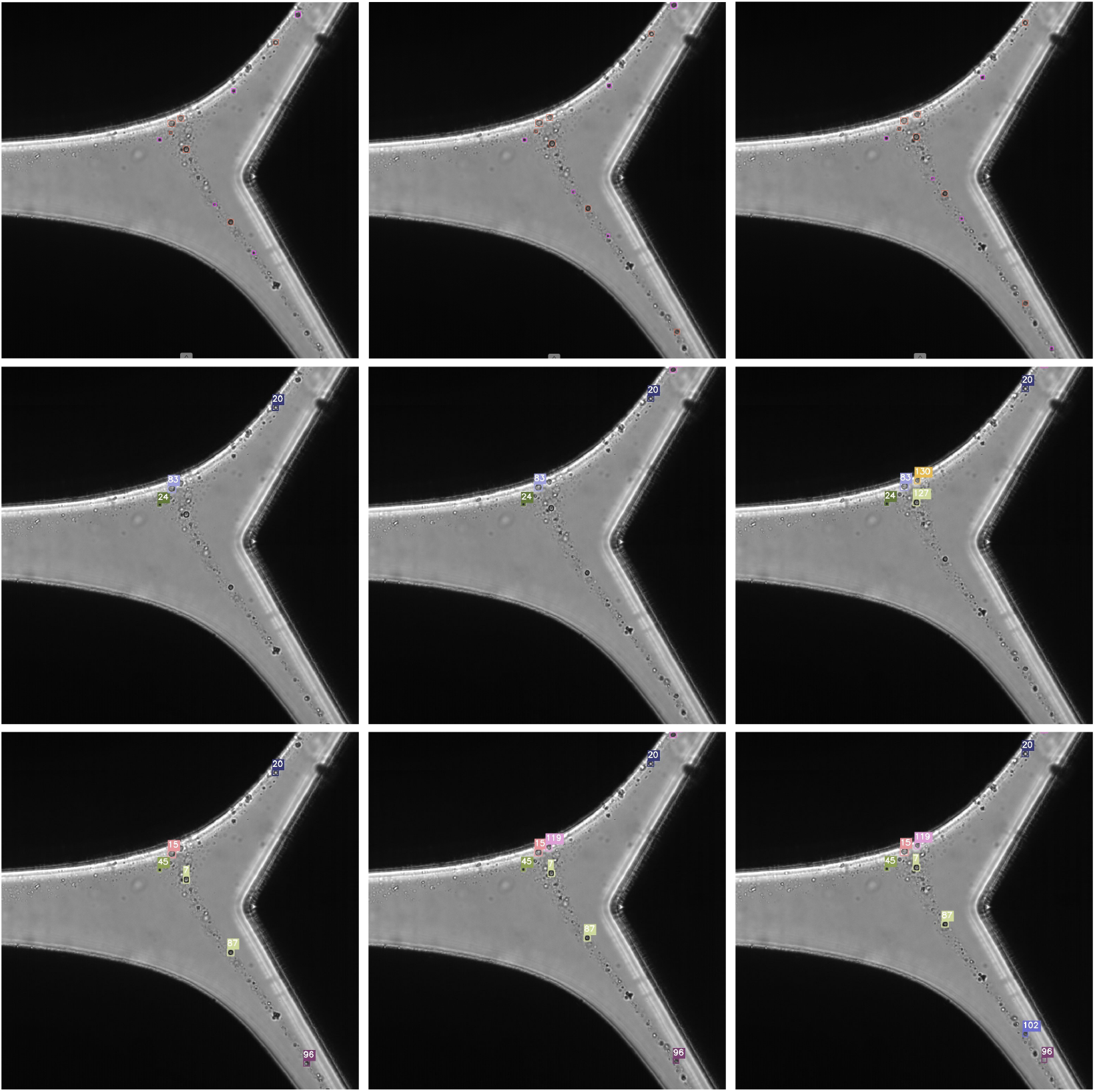
Each row shows three consecutive frames from the testing dataset. The first row is the groundtruth annotation of the cells we are interested in. The second row and the third row are the tracking results from the baseline DeepSORT method, and our proposed method, respectively. The numbers above the bounding boxes indicate the track identifier. The track identifier of the same object is different in the two methods’ results because methods initiated different amounts of tracks before these three frames. We focus on the initialization and perseverance of track identifiers of same objects on each row, then compare two methods. It can be seen that our proposed method picks up more cells as well as starts to track them earlier when compared with the baseline DeepSORT method. A detailed analysis can be found in the discussion portion of the experimental results section.

However, tracking cells in the high velocity area of the chip remains a challenging problem, especially when the cell’s visual appearance is not consistent across the frames. For example, Figure 5 shows the appearance of three cells in different frames, we can see the appearance of melanoma cells in (a) and (b) changes a lot within only 5 frames and the WBC in row (c) has a blurry looking. The appearance feature extraction model used in our method is the one used in DeepSORT, which is trained on a pedestrian dataset. This model may not be able to differentiate individual cells well, and therefore may fail in associating the same cell across frames in this challenging condition. We also notice that most of the lost cells are WBC at the input and exit regions of the chip. The WBC cells are smaller, thus may travel faster and their appearance features are even less distinct, making it more challenging to associate them with the correct tracks.

With approximately 30 MOTA and IDF1 scores, our method still has room for improvements. On the other hand, all three comparison methods achieve over 60 in MOTA on the MOT16 benchmark for general object tracking of pedestrians and cars, but below 20 in our cell tracking problem. Our cell tracking problem is so challenging that existing MOT methods’ performance would drop significantly. By integrating the representative velocity field information into the motion model, we successfully benefit from domain knowledge and improve the cell tracking results.

## VI. Conclusion and Future Work

Our proposed method utilizes both cells’ motion information and appearance features with the help of the prior knowledge of the cells’ velocities for cell tracking works well relative to the challenges of tracking cells in the chip. We use a modified Kalman filter with a non-constant velocity model and the chip’s representative velocity field as an additional input to the Kalman filter to improve the performance of tracking cells in the chip. In the future, we will examine other neural networks to extract appearance features to better distinguish and re-identify the individual cells. As more annotated data becomes available, we will also explore deep learning motion estimation methods and graph network based track association methods to better predict the cell’s complex motion pattern in the chip.

## VII. Acknowledgments

This research was supported by HP Inc. Address all correspondence to Edward J. Delp at

## Footnotes

1 The use of the terms “label” or “labeling” in this paper indicates the process of adding labels such as dyes, dye functionalized probes (e.g., dye functionalized antibody), or particles to the cells, which binds to the cells and alters the cells’ optical properties and enhances their contrast during imaging. It is not to be confused with the ground truthing the images, which in the machine learning community is also sometimes referred to as “label” or “labeling”.

## Notes

### Competing Interest Statement

The authors have declared no competing interest.

